# Ivermectin binds to the allosteric site (site 2) and inhibits allosteric integrin activation by pro-inflammatory cytokines

**DOI:** 10.1101/2025.04.30.651550

**Authors:** Yoko K Takada, Yoshikazu Takada

## Abstract

Ivermectin is known to have anti-inflammatory properties, but the specifics of this action are unknown. We recently showed that multiple pro-inflammatory cytokines (e.g., FGF2, CCL5, CD40L) bind to the allosteric site (site 2) of integrins and activate them. 25-Hydroxycholesterol, a pro-inflammatory lipid mediator, is known to bind to site 2 and induce integrin activation and inflammatory signals (e.g., IL-6 and TNF secretion), suggesting that site 2 is critically involved in inflammation. We showed that TNF, a major pro-inflammatory cytokine, binds to site 2 and induces integrin activation, like other pro-inflammatory cytokines. We recently showed that two anti-inflammatory cytokines (FGF1 and NRG1) bind to site 2 and inhibit integrin activation by inflammatory cytokines. We hypothesized that ivermectin binds to site 2 but inhibits the integrin activation induced by inflammatory cytokines in a mechanism similar to anti-inflammatory cytokines (site 2 antagonists). Consistently, docking simulation predicts that ivermectin binds to site 2. We found that ivermectin suppressed the activation of soluble β3 integrins by multiple pro-inflammatory cytokines (including TNF) in a dose-dependent manner in cell-free conditions, suggesting that ivermectin acts as an antagonist for site 2. This may be a potential mechanism of anti-inflammatory action of ivermectin.

## Introduction

Previous studies showed that multiple inflammatory cytokines (e.g., CX3CL1, CXCL12, CCL5, CD40L, and CD62P) and pro-inflammatory proteins (e.g., sPLA2-IIA) bind to an allosteric ligand binding site of integrins (site 2) and activate integrins (allosteric activation) (1-5) (4, 6, 7) independent of canonical inside-out signaling (8, 9). A previous study showed that 25-hydroxycholesterol, a major pro-inflammatory lipid mediator, binds to site 2, and activates integrins, and induces pro-inflammatory signaling (e.g., IL-6 and TNF secretion) (10). Therefore, it has been proposed that site 2 is involved in pro-inflammatory signaling. Thus, binding of inflammatory cytokines and inflammatory mediators to site 2 may be a common mechanism of inflammatory signaling. Tumor necrosis factor (TNF) is a major pro-inflammatory cytokine, which play a major role in inflammation, cancer and insulin resistance. It is unclear whether TNF binds to site 2, like several other inflammatory cytokines, and induces integrin activation and inflammatory signals. We recently showed that several anti-inflammatory cytokines bind to site 2 (e.g., FGF1 and NRG1) and inhibited integrin activation instead of activating integrins. These findings suggest that FGF1 and NRG1 competing with pro-inflammatory cytokines for binding to site 2 and thereby inhibit site 2-mediated integrin activation and inflammatory signaling (11, 12).

Ivermectin (IVM), a broad-spectrum anthelmintic agent, has anti-inflammatory properties, and affects cellular and humoral immune responses (13). IVM could act through GABA receptors and ligand-gated channels, such as glutamate-gated chloride channels (14). In addition, IVM has been utilized in a wide variety of conditions, from regulating glucose and cholesterol levels in diabetic mice to suppressing cancer proliferation and inhibition of viral replication. IVM-mediated inhibition of inflammatory cytokines production may occur through the suppression of the NF-κB pathway (15). Farnesoid X receptor (FXR) has important roles in maintaining bile acid and cholesterol homeostasis. It has also been reported that ivermectin is a high affinity FXR ligand (16). The mechanism of anti-inflammatory action of IVM, however, has not been established. We hypothesized that anti-inflammatory IVM may bind to site 2 and inhibits the binding of pro-inflammatory cytokines and thereby suppress inflammatory signaling.

In the present study, we describe that docking simulation predicted that IVM binds to site 2 and showed that its binding site overlapped with those of pro-inflammatory cytokines. We showed that docking simulation predicts that TNF binds to site 2 and induces allosteric activation of integrins, like several other inflammatory cytokines, suggesting that pro-inflammatory cytokines induce integrin activation and inflammatory signals. Importantly, IVM inhibited activation of integrins by multiple inflammatory cytokines, including TNF. IVM binding to site 2 may be a potential mechanism of anti-inflammatory action of IVM.

## Results and Discussion

### TNF binds to site 2 and induces integrin activation

Docking simulation predicted that TNF binds to site 2 of integrins (docking energy -19.15 kcal/mol) and activates soluble integrins (Fig. 1). Predicted amino acid residues of αvβ3 for binding to TNF are shown in Table 1. We found that TNF activated soluble integrins αvβ3 and αIIbβ3 in a dose-dependent manner in cell-free conditions. These findings suggest that TNF, like several other pro-inflammatory cytokines, binds to site 2 and activate integrins. It should be noted that TNF-induced activation of soluble integrins is independent of TNF receptors or inside-out signaling, like integrin activation induced by several other pro-inflammatory cytokines (see Introduction).

**Fig. 1.**
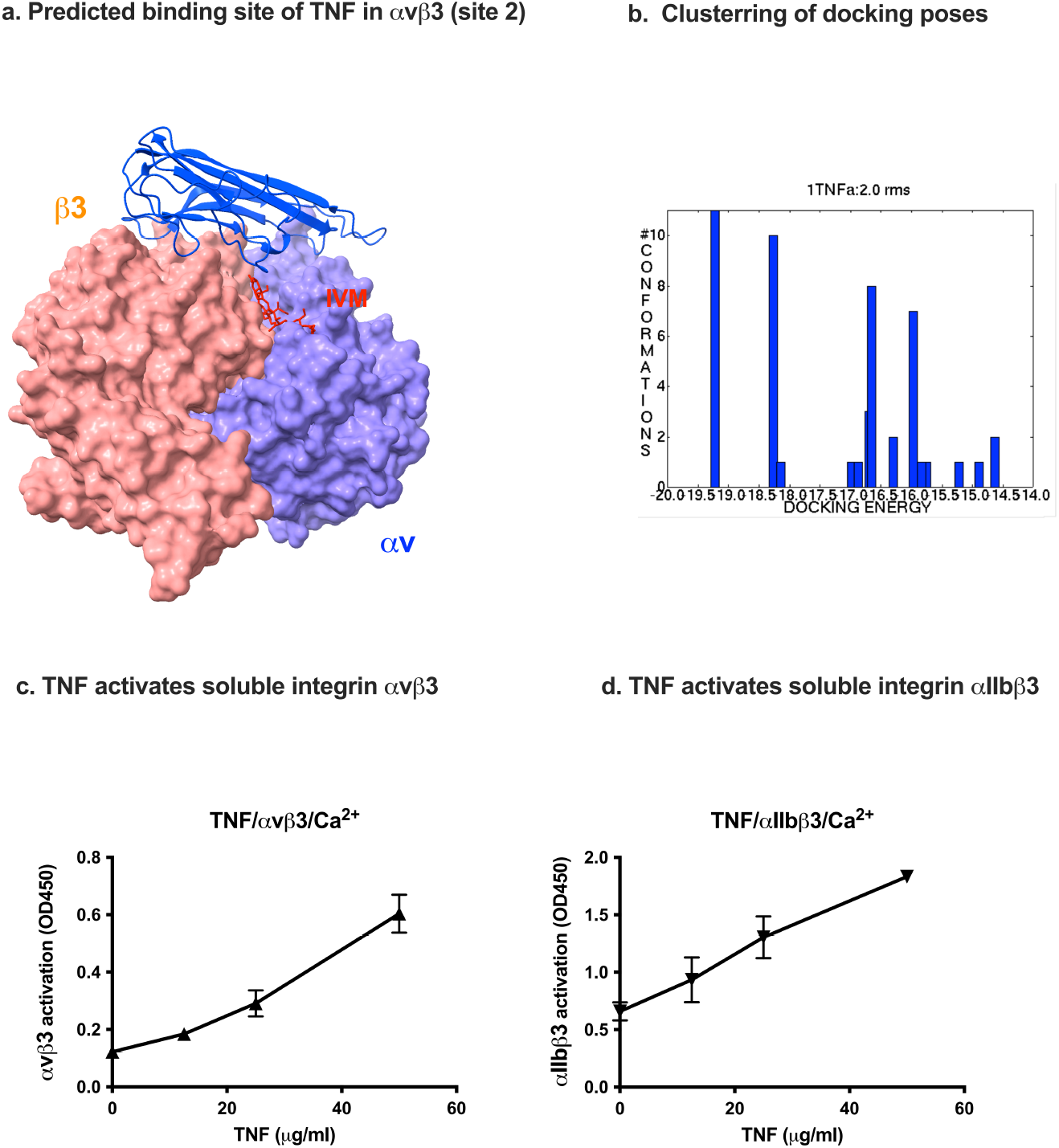
Binding of TNF to site 2 of αvβ3 allosterically activates soluble integrins. (a). Docking simulation between αvβ3 (closed headpiece, 1JV2.pdb) and TNF was performed using autodock3. Position of IVM (red) is shown as a reference (See Fig. 2). IVM and TNF-binding sites overlap. (b). Docking pose of IVM was superposed. The pose in the first cluster is used for further analysis. (c and d). TNF activates integrins αvβ3 (c) and αIIbβ3 (d). Wells of 96-well Immulon 2 microtiter plates were coated with 100 µL PBS containing γC390-411 for αIIbβ3 and γC399tr for αvβ3 for 2 h at 37°C. Soluble recombinant αIIbβ3 or αvβ3 (1 µg/mL) in the presence of TNF was added to the wells and incubated in Hepes–Tyrodes buffer with 1 mM CaCl2 for 1 h at room temperature. Bound αIIbβ3 or αvβ3 was measured using anti-integrin β3 mAb (AV-10) followed by HRP-conjugated goat anti-mouse IgG and peroxidase substrates.

**Table 1.** Predicted amino acid residues involved in binding of TNF to site 2 in integrin αvβ3.

| TNF (1TNF.pdb) | $\alpha\text{v}$ (1JV2) | $\beta 3$ (1JV2) |
| --- | --- | --- |
| Asn19, Glu23, Gln25, Gln27, Trp28, leu29, Arg31, Glu42, Arg44, Asp45, Asn46, Gln47, Val49, Cys69, Pro70, Ser71, Thr72, His73, Leu75, Leu76, Thr77, Thr79, Ser81, Ile83, Val85, Gln88, Thr89, Lys90, Val91, Asn92, Ile97, Gln102, Arg103, Thr105, Glu107, Gly129, Arg131, Glu135, Asn137 | Gly76, Asn77, Asp79, Ala81, Lys82, Asp83, Asp84, Pro85, | Glu171, Glu174, Asn175, Thr182, Thr183, Cys184, Leu185, Pro186, Lys191, His192, Val193, Leu194, Thr195, Arg202, Glu205, Glu206, Val207, Lys208, Lys209, Gln210, Ser211, Gly276, Ser277, Asp278, Asn279, His280, Ser282, Thr285, Thr286 |

Amino acid residues within 0.6 nm between ivermectin and $\alpha\beta 3$ were selected using Pdb Viewer (version 4.1).

**Table 2.**
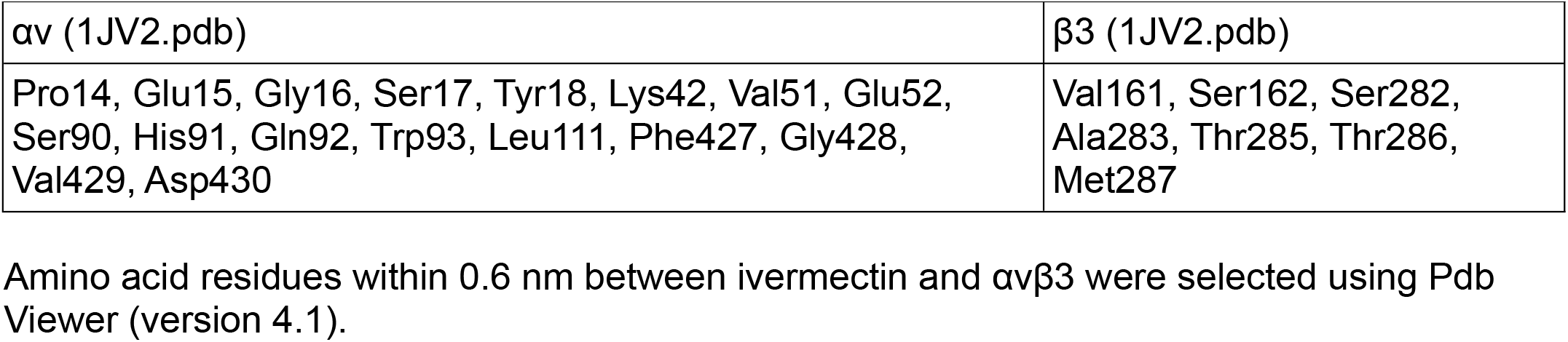
Predicted amino acid residues involved in binding to ivermectin in integrin αvβ3.

### Docking simulation predicts that IVM binds to site 2 of integrin αvβ3

Previous studies showed that anti-inflammatory cytokines FGF1 and NRG1 (site 2 antagonists) inhibit the binding of pro-inflammatory cytokines (site 2 agonists) and subsequent integrin activation (see Introduction). To address this hypothesis, we performed docking simulation of interaction between integrin αvβ3 (closed headpiece, 1JV2.pdb) and IVM. The simulation predicted that IVM binds to site 2 (docking energy -18.24 kcal/mol). (Fig. 2a). Predicted amino acid residues involved in IVM binding are shown in Table 1. (Fig. 2b).

**Fig. 2.**
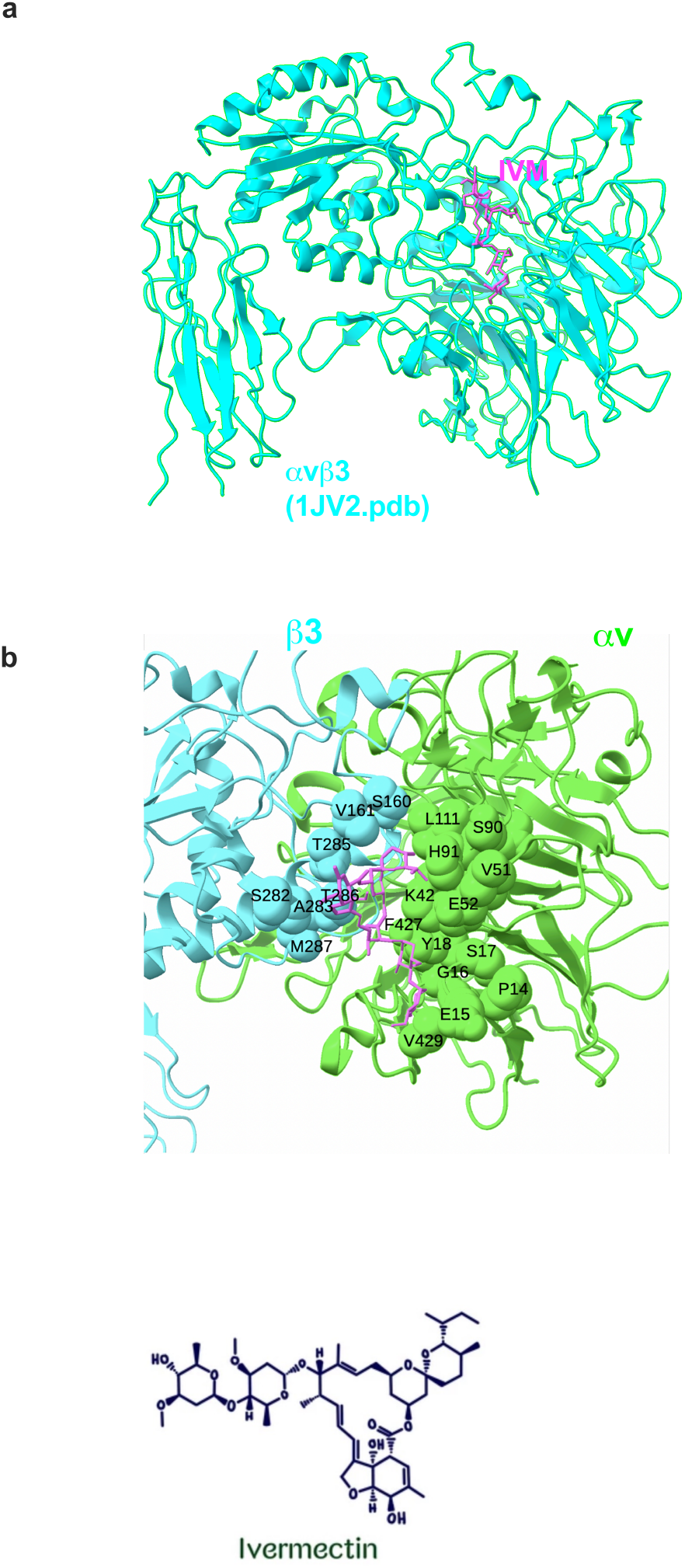
Docking simulation of interaction between integrin αvβ3 and ivermectin (IVM). (a) Docking simulation between αvβ3 (closed headpiece, 1JV2.pdb) and IVM was performed using autodock3. Docking energy is -18.24 kcal/mol. (b) Amino acid residues of αvβ3 surrounding IVM in site 2.

The predicted IVM-binding site is in the center of site 2 and overlaps with those of multiple pro-inflammatory cytokines, suggesting that IVM may inhibit integrin activation by pro-inflammatory cytokines.

### IVM inhibits allosteric integrin activation induced by multiple pro-inflammatory cytokines

We studied if IVM inhibits integrin activation by multiple pro-inflammatory cytokines. IVM suppressed activation αvβ3 (Fig. 3a-d) and αIIbβ3 (Fig. 4 a-d) by FGF2, CD40L, CCL5 and TNF, suggesting that IVM acts as an antagonist for site 2 in αvβ3 and αIIbβ3. This is consistent with the idea that anti-inflammatory action of IVM is mediated by blocking the binding of multiple inflammatory cytokines to site 2.

**Fig. 3.**
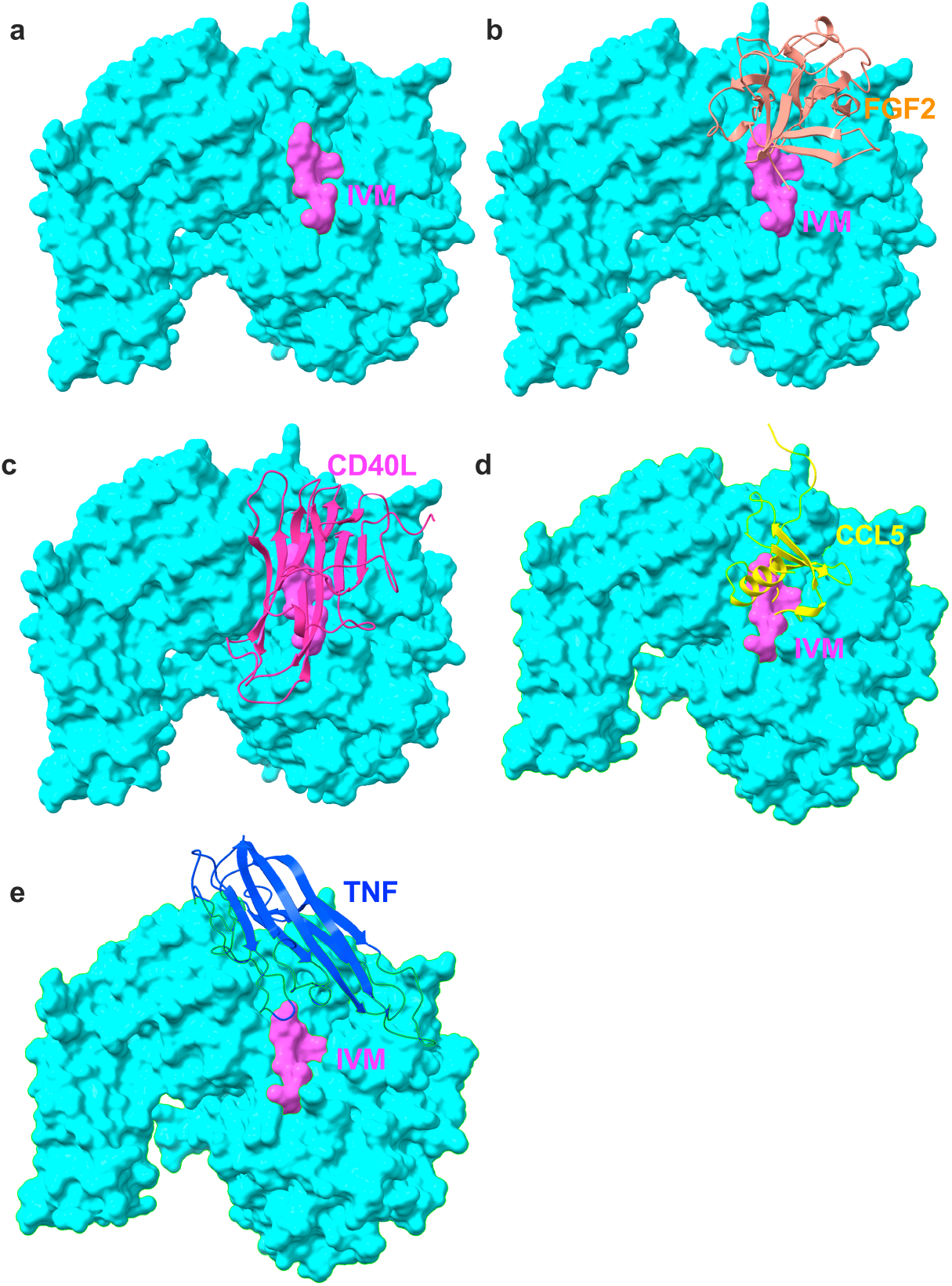
Positions of IVM and inflammatory cytokines on site 2 of αvβ3. IVM and multiple inflammatory cytokines that bind to site 2 were superposed. Docking poses of inflammatory cytokines were taken from previous papers. (a) IVM only, (b) FGF2 (11) and IVM, (c) CD40L (6) and IVM, (d) CCL5 (5) and IVM, and (e) TNF (docking energy -19.15 kcal/mol, the present study) and IVM.

**Fig. 4.**
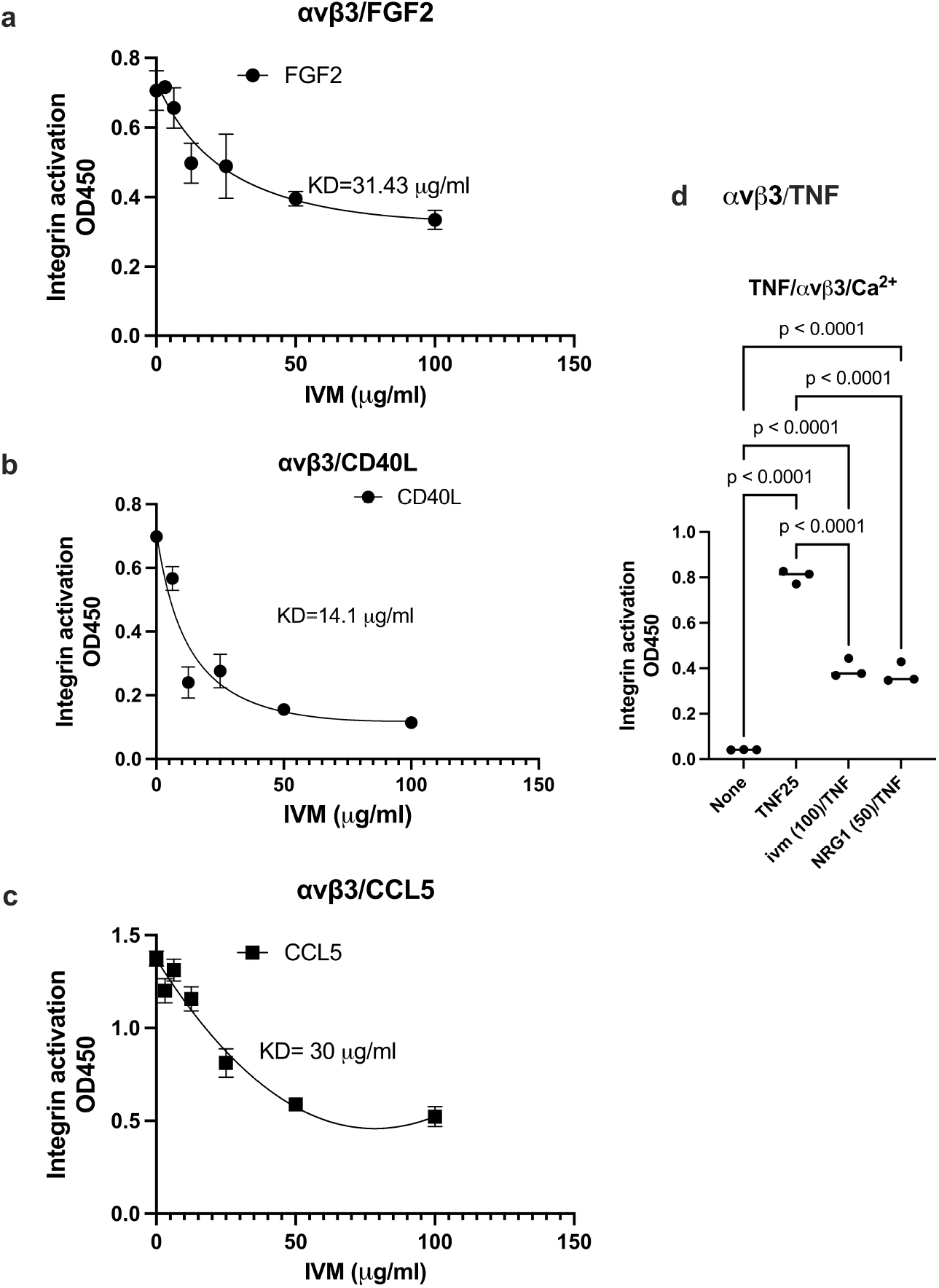
Inhibition by IVM of pro-inflammatory cytokine-mediated activation of integrin αvβ3. Wells of 96-well microtiter plate were coated with the C-terminal domain of fibrinogen γ-chain in which the C-terminal 12 residues are truncated (γC399tr), a specific ligand for integrin αvβ3 (50 µg/ml). Soluble αvβ3 was activated by (a) FGF2 (12.5 µg/ml), (b) CD40L (25 µg/ml), (c) CCL5 (at 6.25 µg/ml) in the presence of IVM. (d) TNF (25 µg/ml). NRG1 (50 µg/ml), and IVM (100 µg/ml). Data is shown as means +/- SD in triplicate experiments.

We showed that TNF-induced activation of integrins is also inhibited by IVM (Fig. 5). These findings suggest that IVM suppress insulin resistance by inhibiting TNF-induced integrin activation.

**Fig. 5.**
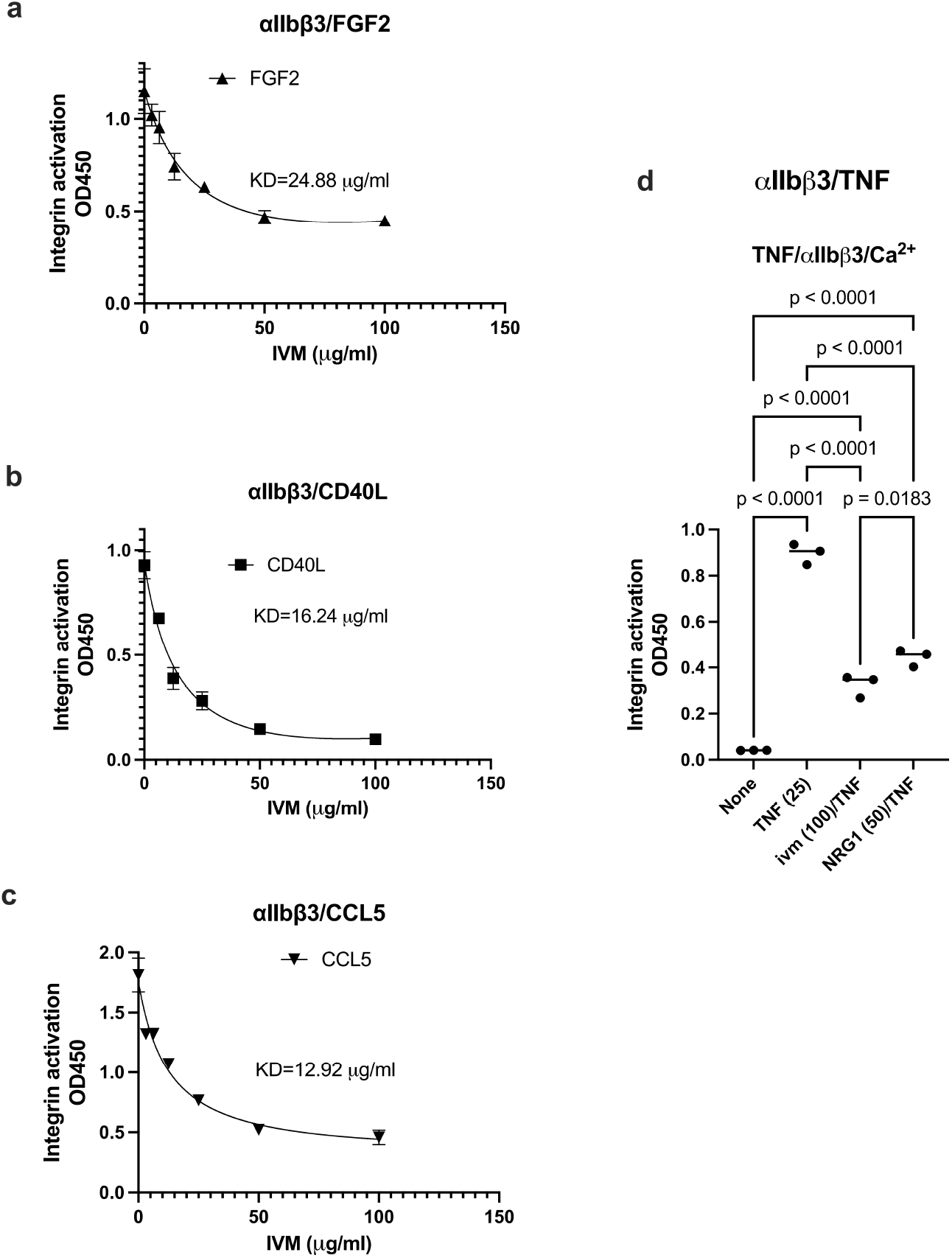
Inhibition by IVM of pro-inflammatory cytokine-mediated activation of integrin αIIbβ3 Inhibition by IVM of pro-inflammatory cytokine-mediated activation of integrin αIIbvβ3. Wells of 96-well microtiter plate were coated with the C-terminal domain of fibrinogen γ-chain C-terminal 12 residues (γC390-411), a specific ligand for integrin αIIbβ3 (20 µg/ml). Soluble αIIbβ3 (1 µg/ml) was activated by (a) FGF2 (12.5 µg/ml), (b) CD40L (25 µg/ml), (c) CCL5 (at 6.25 µg/ml) in the presence of IVM. (d) TNF (25 µg/ml). NRG1 (50 µg/ml), and IVM (100 µg/ ml). Data is shown as means +/- SD in triplicate experiments.

### The present findings will facilitate the study of IVM as anti-inflammatory, anti-cancer, anti-virus and anti-thrombotic action

IVM has been widely used in clinic originally as an anti-parasite agent. It has been proposed that IVM is anti-inflammatory, anti-cancer and anti-virus activity as well. However, IVM has not been systematically tested in clinic if IVM is effective as anti-inflammatory, anti-cancer and anti-viral agent. The present study provides evidence for the first time that IVM is an inhibitor of site 2-mediated integrin activation (and subsequent inflammatory signaling) (Fig. 6). Allosteric activation of αvβ3 by inflammatory cytokines may be related to pro-inflammatory action of inflammatory cytokines. Also, inflammation is involved in cancer proliferation and multiple pro-inflammatory cytokines (e.g., CCL5, CX3CL1, CXCL12, and TNF) play a role in cancer. Blocking pro-inflammatory signals through site 2 may suppress cancer proliferation. Activation of platelet integrin αIIbβ3 is a key event in platelet activation and thrombus formation (8, 17). We previously showed that multiple cytokines bind to site 2 of αIIbβ3 and activated this integrin (5). It is highly likely that multiple inflammatory cytokines allosterically activate αIIbβ3 and induce platelet activation. IVM may thus have anti-thrombotic action (e.g., deep vein thrombosis). The present study is expected to facilitate the elucidation of the mechanism of action of IVM and repurposing of IVM.

**Fig. 6.**
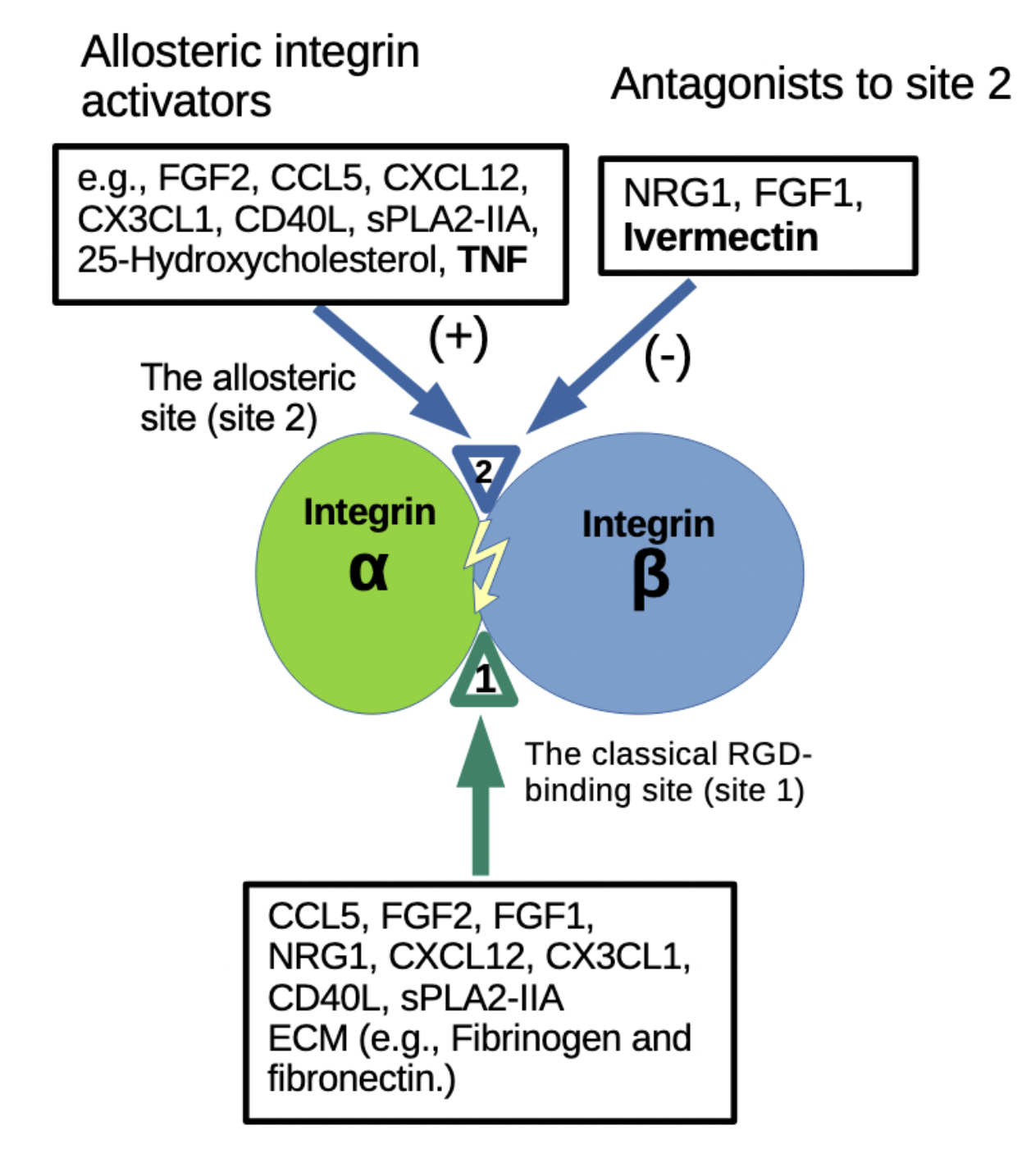
A model of inhibition of binding of pro-inflammatory cytokines to the allosteric site 2 by anti-inflammatory cytokines and ivermectin. Previous studies showed that multiple pro-inflammatory cytokines and 25-hydoxycholesterol bind to the center of site 2 and induce integrin activation and inflammatory signals (see Introduction). TNF also binds to site 2 and induces integrin activation (the present study). In contrast, FGF1 and NRG1 bind to site 2 (off the center of site 2) and inhibit integrin activation by inflammatory cytokines (see Introduction). Ivermectin binds to the center of site 2 and inhibits binding of pro-inflammatory cytokines and resulting integrin activation and inflammatory signals. These findings suggest that ivermectin binding to site 2 is a potential mechanism of anti-inflammatory action of ivermectin.

IVM is known to be insoluble to water (max. 5 µg/ml) and we used stock IVM (10 mM) in 95% ethanol and diluted in water. Although we showed that calculated 100 µM IVM effectively inhibited allosteric activation of integrins, actual IVM concentration may not be higher than 5 µg/ml (approx. 5 µM). If the binding of IVM to site 2 is critical for its anti-inflammatory action, it would be possible to design more effective or water soluble IVM variants.

## Materials and methods

### Materials

#### Protein Expression

The truncated fibrinogen γ-chain C-terminal domain (γC399tr) was generated as previously described (18). Fibrinogen γ-chain C-terminal residues 390-411 cDNA encoding (6 His tagged) [HHHHHH]NRLTIGEGQQHHLGGAKQAGDV] was conjugated with the C-terminus of GST (designated γC390-411) in pGEXT2 vector (BamHI/EcoRI site). The protein was synthesized in E. coli BL21 and purified using glutathione affinity chromatography. FGF2 was synthesized as previously described (19). CCL5 (5), CXCL12 (3), CX3CL1 (20) were synthesized as described. TNF was synthesized as described for CCL5 and CX3CL1. Briefly, cDNA for TNF [VRSSSRTPSDKPVAHVVANPQAEGQLQWLNRRANALLANGVELRDNQLVVPSEGLYLIYSQ VLFKGQGCPSTHVLLTHTISRIAVSYQTKVNLLSAIKSPCQRETPEGAEAKPWYEPIYLGGVF QLEKGDRLSAEINRPDYLDFAESGQVYFGIIAL] was synthesized and subcloned into the BamH1/EcoRI site of PET28a vector. Protein was synthesized in E. coli BL21 (endotoxin-free, Clea coli) as insoluble inclusion bodies. TNF was purified by in denaturing conditions in 8M urea in Ni-NTA affinity chromatography and refolded as described (20).

#### Docking simulation

Docking simulation of the interaction between TNF/IVM and integrin αvβ3 (closed head-piece form, PDB code 1JV2) was performed using AutoDock3, as described previously (21). We used the headpiece (residues 1–438 of αv and residues 55–432 of β3) of αvβ3 (closed form, 1JV2.pdb). Cations were not present in integrins during docking simulation.

### Statistical Analysis

Treatment differences were tested using ANOVA and Tukey multiple comparison tests to control the global type I error using Prism 10 (Graphpad Software, Boston, MA, USA).

Activation of soluble αIIbβ3 and αvβ3 by ELISA-type activation assays were performed as described previously (2, 20). Briefly, wells of 96-well Immulon 2 microtiter plates (Dynatech Laboratories, Chantilly, VA, USA) were coated with 100 µL 0.1 M PBS containing γC390-411 for αIIbβ3 and γC399tr for αvβ3 for 2 h at 37°C. The remaining protein-binding sites were blocked by incubating with PBS/0.1% BSA for 30 min at room temperature. After washing with PBS, soluble recombinant αIIbβ3 or αvβ3 (1 µg/mL) in the presence or absence of FGF1 and/or FGF2 was added to the wells and incubated in Hepes–Tyrodes buffer (10 mM HEPES, 150 mM NaCl, 12 mM NaHCO_3_, 0.4 mM NaH_2_PO_4_, 2.5 mM KCl, 0.1% glucose, 0.1% BSA) with 1 mM CaCl_2_ for 1 h at room temperature. After unbound αIIbβ3 or αvβ3 was removed by rinsing the wells with binding buffer, bound αIIbβ3 or αvβ3 was measured using anti-integrin β3 mAb (AV-10) followed by HRP-conjugated goat anti-mouse IgG and peroxidase substrates.

## Acknowledgement

This work is partly supported by the UC Davis Comprehensive Cancer Center Support Grant (CCSG) awarded by the National Cancer Institute (NCI P30CA093373). The contents reported/presented within do not represent the views of the Department of Veterans Affairs or the United States Government.

## Notes

### Competing Interest Statement

The authors have declared no competing interest.

